# Predicting Methylation from Sequence and Gene Expression Using Deep Learning with Attention

**DOI:** 10.1101/491357

**Authors:** Alona Levy-Jurgenson, Xavier Tekpli, Vessela N. Kristensen, Zohar Yakhini

## Abstract

DNA methylation has been extensively linked to alterations in gene expression, playing a key role in the manifestation of multiple diseases, most notably cancer. For this reason, researchers have long been measuring DNA methylation in living organisms. The relationship between methylation and expression, and between methylation in different genomic regions is of great theoretical interest from a molecular biology perspective. Therefore, several models have been suggested to support the prediction of methylation status in samples. These models, however, have two main limitations: (a) they heavily rely on partially measured methylation levels as input, somewhat defeating the object as one is required to collect measurements from the sample of interest before applying the model; and (b) they are largely based on human mediated feature engineering, thus preventing the model from unveiling its own representations. To address these limitations we used deep learning, with an attention mechanism, to produce a general model that predicts DNA methylation for a given sample in any CpG position based solely on the sample's gene expression profile and the sequence surrounding the CpG.

We show that our model is capable of generalizing to a completely separate test set of CpG positions and subjects. Depending on gene-CpG proximity conditions, our model can attain a Spearman correlation of up to 0.8 and MAE of 0.14 for thousands of CpG sites in the test data. We also identify and analyze several motifs and genes that our model suggests may be linked to methylation activity, such as Nodal and Hand1. Moreover, our approach, and most notably the use of attention mechanisms, offers a novel framework with which to extract valuable insights from gene expression data when combined with sequence information.

The code and trained models are available at: https://github.com/YakhiniGroup/Methylation

## I. INTRODUCTION

DNA methylation is a chemical process that modifies DNA in living organisms and can significantly affect gene expression, mostly through the inhibition of transcription. In humans, DNA methylation refers to the presence of a methyl group at a defined position of a cytosine and occurs mostly in CpG dinucleotides. It has been particularly shown to affect gene expression in gene promoter regions with relatively dense CpGs, known as CpG islands (CGI). When a large number of proximal CpGs are methylated, the transcription of nearby downstream genes may be inhibited. This process is, for example, prominent in the silent X-chromosome in females [37].

DNA methylation plays a key role in disease development. Specifically, hypermethylation can lead to stable silencing of tumor suppressor genes [21]. This process has therefore been extensively observed and studied in the context of cancer [24], [15], [41], [29]. While various forms of cancer are central to the discussion on DNA methylation, it has also been linked to other diseases and biological processes such as cardiovascular disease [13] and Alzheimer’s Disease [19] as well as gene expression regulation in general [40] and epigenetic editing [35]. Hence, researchers have long been measuring DNA methylation levels.

Currently, there are several methods available for measuring DNA methylation [23]. Some of these methods, however, require specialized protocols or a relatively large DNA sample size. Hence, depending on the required task, the costs could be significant and the data collection may be impractical. For this reason, the prediction of DNA methylation levels through other means could prove highly useful. In addition, and perhaps more importantly, the link between gene expression and methylation is still an open-ended question and predictive analyses may provide insight into this relationship. In this work we set out to address both aspects of methylation.

The contribution of this work includes: (1) We provide a practical tool that enables potential users to input any CpG position for which DNA methylation was not measured, along with the sample’s gene-expression profile, and obtain a prediction. We do so by training a machine learning model that combines gene expression data with the ambient DNA sequence at the CpG of interest. We demonstrate that this model is capable of generalizing across CpG sites as well as across samples. (2) From a theoretical perspective, this result provides proof for a sharp, albeit not necessarily causal, link between sequence and expression and between local methylation events. Furthermore, we observe better predictability for CpGs that reside closer to genes. We also unveil motifs and genes that the model identified as significant contributors to the prediction. Specifically, we link HAND1 and NODAL to methylation activity in the cohort analyzed. (3) We provide a novel model design and framework that support the combination of gene-expression data with genomic sequences to extract valuable molecular-level insights.

### A. Related Work

Over the past decade, researchers have been investigating the use of machine learning for the prediction of methylation. In [4] and [10] the authors used classifiers such as Support Vector Machines (SVMs) and decision trees to determine the status of a given CpG using both structural and sequence-specific features. Similarly, [44] suggested a random forest classifier that uses features such as genomic position and neighbor methylation levels. The latter were noted as significant contributors but clearly require collecting partial methylation data. Others [26], have used a regression approach to predict continuous methylation levels across tissues, also using SVMs. While the use of regression is indeed more appropriate in the context of continous methylation measurements, this approach requires extensive data collection from a source tissue. More recently, [42] used a deep learning model to predict whether a CpG was hypo- or hyper-methylated by using DNA patterns and topological features. The latter are human engineered features taken as input by the network model. Like previous methods, this model is limited to binary classification, and is specifically constrained to hypo-/hyper- methylation.

To conclude, the main limitations posed by previous models include: (1) The need to measure methylation in some (or all, in the case of learning between tissues) CpG sites. (2) Extensive use of human-engineered features. This not only incorporates human biases, but also prevents the model from unveiling novel representations. (3) The majority are binary classifiers when in reality methylation levels are measured continuously, representing fractions of cells with any given status.

## II. Approach

To address the limitations posed by the aforementioned methods, we suggest a general deep learning model that does not require measuring methylation levels in the sample of interest, is not limited to specific CpGs, uses neural networks as feature extractors instead of human-engineered features and provides continuous predictions. Specifically, we predict methylation levels at a given CpG in a given sample based on three factors:

- The sequence surrounding the CpG.
- The sample’s gene expression profile.
- The distance between the profiled genes and the CpG.

We use a generalized approach that can be applied to any set of CpGs. We avoid incorporating human-engineered features derived from the DNA sequence by using a Convolutional Neural Network (CNN) as a motif detector. We also do not manually select genes to include in the model, but rather incorporate the gene-expression profile by using three attention mechanisms that take the input context into consideration. That is - the attention mechanism is determined by the sequence around the CpG of interest, the distance between the CpG of interest to each of the genes and the gene-expression profile as a whole (see Figure 2). We test our model on a completely separate set of CpG positions and subjects to ensure that our model can indeed generalize well. Our method is implemented in TensorFlow for Python, and is available online^1^.

## III. METHODS

### A. Datasets

We used data from two cancer cohorts: (1) 782 breast cancer patients and (2) 498 prostate cancer patients. For each patient, we obtained two types of data: (a) gene expression data in RSEM normalized count for 17,997 genes (RNA-seq) and (b) methylation levels at 360,531 CpG sites (450K Illumina array).

In addition to the patient-specific data, we also use data specific to any CpG locus: (a) the ambient sequence - 399 base-pairs upstream of the CpG and 399 downstream, for a total of 800 base-pairs and (b) the genomic distance between each gene in the profile considered and the locus of interest.

### B. Constructing the Model

Our task is to predict the methylation level at a CpG site in a sample taken from a given subject, using the samples’s gene expression profile and the ambient sequence at the CpG site. To do so, we created a multi-modal neural network comprised of four sub-networks: one CNN, which acts as a motif detector for the surrounding sequence, and three attention components which act as gene amplifiers, each based on the input provided. These sub-networks are then combined into a single fully-connected network to produce the final prediction.

#### Input Data

We will define a single training example to represent one subject (or sample) and one CpG. It contains the following components:

1. The subject’s gene expression vector ***e***, where each entry, ***e_i_***, represents the expression level of a gene ***g_i_***.
2. The sequence surrounding the CpG of interest, represented as a one-hot matrix ***S***.
3. A vector ***d***, where ***d_i_*** is computed based on the distance, in base-pairs, between ***g_i_*** and the CpG of interest. Specifically, a gene residing within the first 2,000 base-pairs received a value of 1, the next 2,000 a value of 0.5 and so on until the last bucket of 2,000 was given a value of 0.5^9^. Beyond this point ***d_i_*** was set to 0. For genes residing on a different chromosome this value was also set to 0.

The leftmost layers (orange) in Figure 2 illustrate a single training sample.

#### Convolutional Neural Network as a Motif Detector

As seen in previous work [1], [3], we used a CNN as a motif detector. The CNN contains several filters that scan through the sequence and identify motifs of interest. More formally, given an input sequence *seq*, we convert it into a matrix ***S*** such that each row is a one-hot vector representing a single nucleotide base out of the five options: A,C,G,T,N. Multiple filters of size 11x5 convolve over ***S***, followed by batch normalization, a non-linearity layer and a pooling layer. This last layer allows for small shifts in the motif’s position between sequences. The pooling layer is then followed by 3 fully-connected layers that result in a vectorized representation ***s*** of the original sequence *seq*.

#### Attention Mechanism for Gene-Expression

To incorporate the gene-expression profile, we have used attention mechanisms [17]. An attention mechanism is essentially a vector of probabilities usually obtained by employing softmax on the final output layer of a neural network. This vector in turn is used as a filter for another vector, often via an element-wise product. In our case, we created three attention vectors, each of which is derived from the output vector of a different neural network. We then multiplied each of them element-wise by the gene-expression profile vector e as seen in Figure 2. Specifically, we created the following three neural networks to generate three attention vectors:

1. A second CNN that operates on ***S*** as above, with output layer ***a_seq_***.
2. A fully-connected neural network based on the distance vector ***d*** with output layer ***a_dist_***.
3. A fully-connected neural network based on the gene-expression vector ***e*** with output layer ***a_exp_***.

These attention mechanisms enable the model to select which genes are important given any input context and provide a form of conditional importance to all measured expression levels. For example, the first attention vector might detect the presence of a transcription factor binding site (TFBS) proximal to the CpG via motifs learned by the CNN. That transcription factor can be related to methylation activity (e.g. the transcription factor HAND1 plays a key role in the development and differentiation of cell lineages during embryonic development [2] - a process which is also known to be largely governed by methylation activity [28]). Hence, its expression levels might affect methylation status around its binding sites (but not necessarily otherwise). The second attention mechanism might learn that a gene that is in close proximity to the given CpG has higher predictive value than a gene residing on a separate chromosome. The third might detect that a certain combination of co-expressed genes is informative of the expression level of some other gene.

Each of the three attention vectors is multiplied elementwise by the gene-expression profile ***e*** to produce three vectors of the following form:

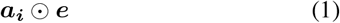

where ⊙ is the element-wise product and ***a_i_*** is one of the three attention vectors.

#### The Combined Multi-Modal Neural Network

Having described how the sequence was processed using a convolutional neural network, as well as the attention mechanisms employed on the gene-expression profile, we are ready to combine the two components into a single neural network. Similar to previous work [17], we combined the output layers of both the CNN and the attention mechanism via concatenation and fed the concatenated representation into a final fully-connected neural network. More formally, denoting the output layer representing the surrounding sequence as ***s***, we form the following input to the final fully-connected network:

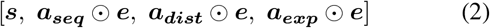

The full network architecture is described in Figure 2.

### C. Training

#### Training Tasks

We create a unified model by training three different submodels designed to address three close, but not identical, prediction tasks. One focuses on CpGs with a gene that is in close proximity, specifically within a window of 2,000 base-pairs on either side. The second focuses on CpGs with a gene that is in medium-proximity of 10,000 base-pairs, and the third is applicable to all CpGs regardless of gene proximity. To ensure that our model learns to generalize across different samples, we first created a unified dataset that includes both the breast cancer and prostate cancer cohorts, as described in Section III-A. Then, for each task we created a new dataset that matched the task's criteria. For the first two tasks we took all CpGs from the combined dataset that satisfied the relevant window criteria, leaving us with 9,563 and 73,828 CpGs respectively. For the third task we randomly sampled 99,981 CpGs out of the total 360K available. We then randomly split each of the resulting three datasets into training, validation and test sets (see Table I for the exact breakdown). The purpose of generating these three sub-models is twofold: (1) it enables us to provide more accurate predictions under certain proximity conditions and (2) having a model specialize solely on proximity data (as opposed to generic CpG data), especially in the case of Model 2 where many CpGs satisfy the 10,000 window condition, will enable it to learn more effectively those gene-CpG representations we sought out to discover, further facilitating our analysis and interpretation of the resulting representations.

**TABLE I.**
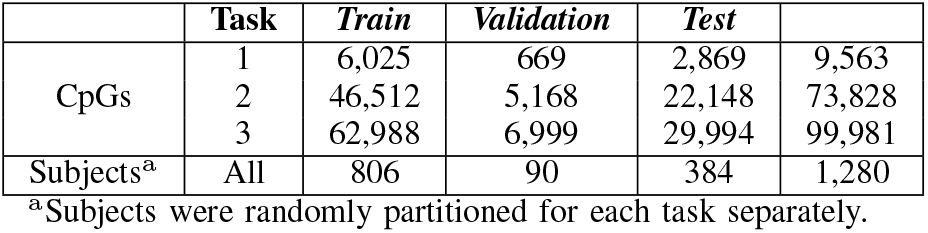
Data Breakdown for Training and Evaluation.

It is important to note that in all cases, the validation and test sets contain only subjects and CpGs that did not participate in the training phase at all (as described in Figure 1). The purpose of this was to evaluate the extent to which the model is capable of generalizing beyond previously seen subjects as well as beyond previously seen CpGs.

**Fig. 1.**
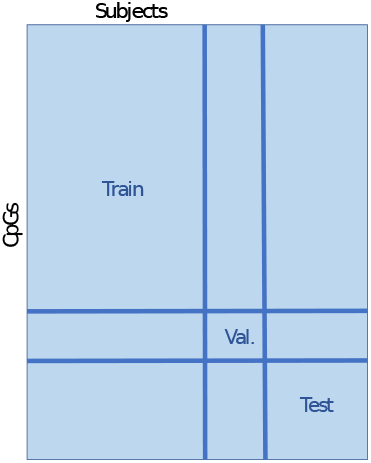
Train vs test set. The test set consists solely of subjects/samples (columns) and CpGs (rows) that have not been included in the train or validation set. The association of each subject and CpG to either one of train, validation or test sets was random. The test set, therefore, includes samples from both cancer types.

#### Training Specifications

We have tested various network configurations for each of our models, with depths ranging from 3 to 5 layers per subnetwork and hidden neurons numbered between 50 to 500. The exact number of layers and parameters for each part of the network used in all subsequent steps appear in Figure 2.

**Fig. 2.**
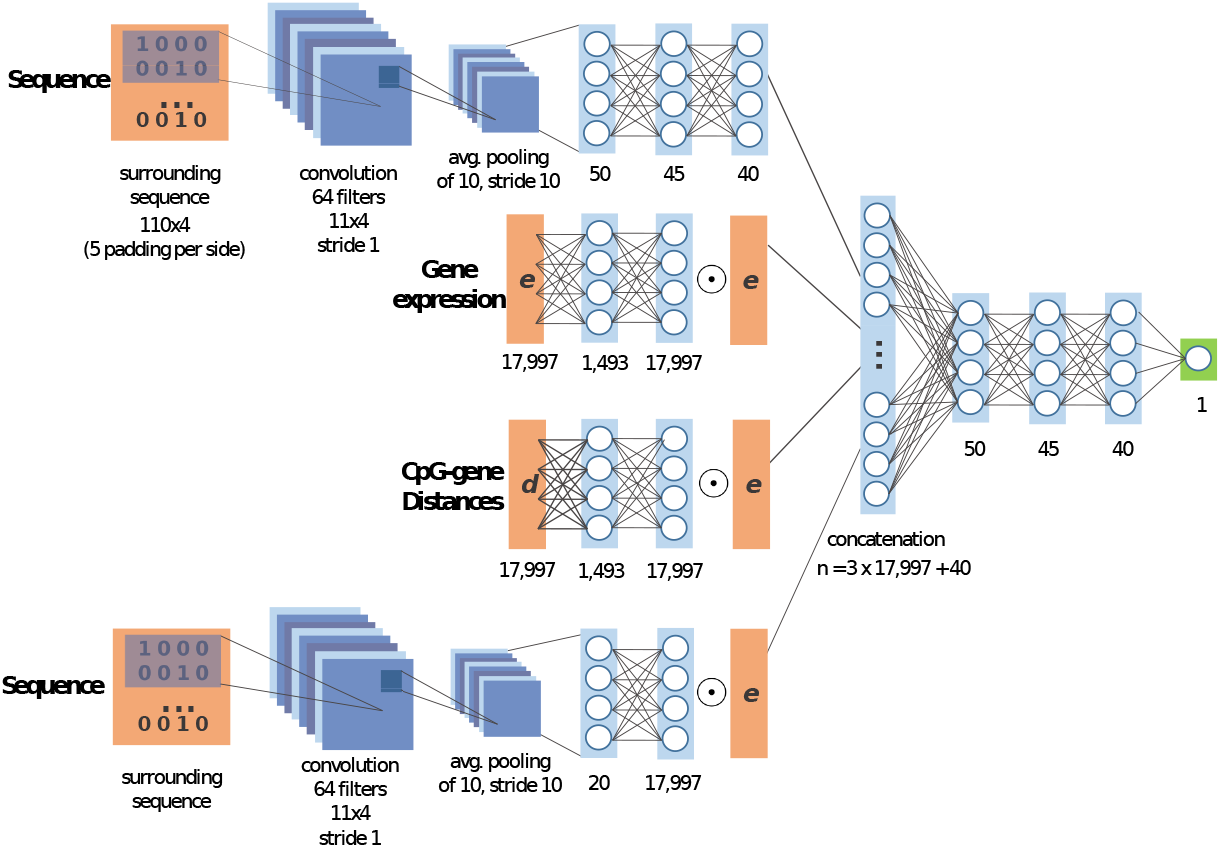
The full model. A multi-modal neural network is constructed to combine the CNN with the three attention components. From top to bottom: the CNN motif detector, the gene-expression based attention mechanism, the distance-based attention mechanism and the sequence-based attention mechanism. The symbol ⊙ stands for the element-wise product. The final layers of these sub-networks are combined via concatenation into a single vector which is then fed into a final neural network to allow for multi-modal representations. The input layers appear in orange, hidden layers are in blue and the final output layer, representing the predicted level of methylation at the CpG of interest and in the sample of interest, is in green.

Our models were trained using the Adam Optimizer [22] with an initial learning rate of 0.01 and a mini-batch size of 300. We applied batch normalization to each layer, except for the output layer, followed by an ELU non-linearity [6]. We use the mean-absolute error (MAE) as the loss function. Training to convergence on the validation set took roughly 4 hours on a Tesla K80 GPU.

## IV. RESULT

### Predicting Methylation Levels

Recall that our unified model includes three different setups:

1. Model 1 - trained and tested using all available CpGs that are within 2,000 base-pairs to the nearest gene.
2. Model 2 - trained and tested using all available CpGs that are within 10,000 base-pairs to the nearest gene.
3. Model 3 - trained and tested using randomly selected CpGs, regardless of gene proximity.

We evaluated each model on its respective held-out test sets, as described earlier and in Figure 1. Model 1, for gene-proximal CpGs, achieves an MAE of 0.14 and 0.8 Spearman correlation (− log (*p*) > 100) on its held-out test set (see Table I). Model 2, for gene-neighboring CpGs, achieves an MAE of 0.16 and 0.75 Spearman correlation (− log (*p*) > 100) on its held-out test set. Model 3, trained on random, general, CpGs, attains 0.2 MAE and 0.65 Spearman correlation (− log (*p*) > 100). Figure 3 displays the scatter plot of the predicted vs the actual methylation values for Model 2. These results demonstrate that the model better utilizes the attention mechanisms when provided with relevant data. Furthermore, these results also show that the models are capable of generalizing to new CpGs and subjects.

**Fig. 3.**
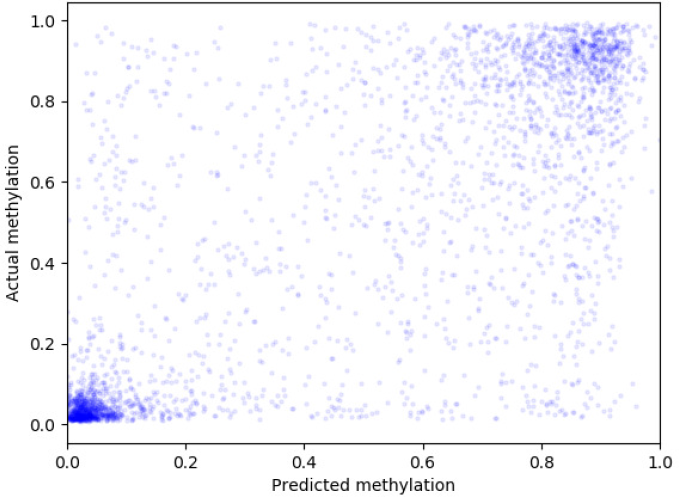
Scatter plot comparing the predicted methylation values of Model 2 and the actual methylation values. We depict 3,000 randomly selected instances from the held-out test set.

### Gene-Expression Attention Learned by the Model

In order to understand the importance attributed to the different genes by the model, we analyzed the output of the attention mechanisms, i.e. the output of the softmax layers. For this purpose we used Model 2.

#### Sequence-based attention

In this section, we set out to understand which genes played an active role in the prediction of methylation at multiple CpG sites. Providing we identify such highly relevant genes, we could further analyze the ambient sequences of the corresponding CpGs as a group, and possibly discover enriched motifs. The first step is to identify genes that were relevant across multiple CpGs. To do so, we gathered all attention vector outcomes for each of the unique sequences in the test set (each representing one CpG site). This is accomplished by feeding the trained sequence-based attention component with one ambient sequence at a time, resulting in a single attention vector per CpG. Recall that each entry in this attention vector corresponds to one gene. Hence, we can slice across CpGs and gather all attention scores attributed to a gene (from all CpGs), resulting in one vector per gene. Because these attention scores can be interpreted as the importance attributed to the gene by the CpG that produced it, we sort each of the gene’s attention values in descending order so that the top of its list contains those CpGs for which the gene received the highest attention score (importance). We then remove all genes that did not have at least one attention score > 0.1. The top 100 CpGs from each of the remaining sorted lists are the columns seen in Figure 4 (a). Notice that the darker cells in each column correspond to CpGs that associated the respective gene with a higher relevance to its prediction (by giving it a higher attention value). Also note that each gene was sorted separately, hence the top CpGs in one gene’s column may differ from those of another.

**Fig. 4.**
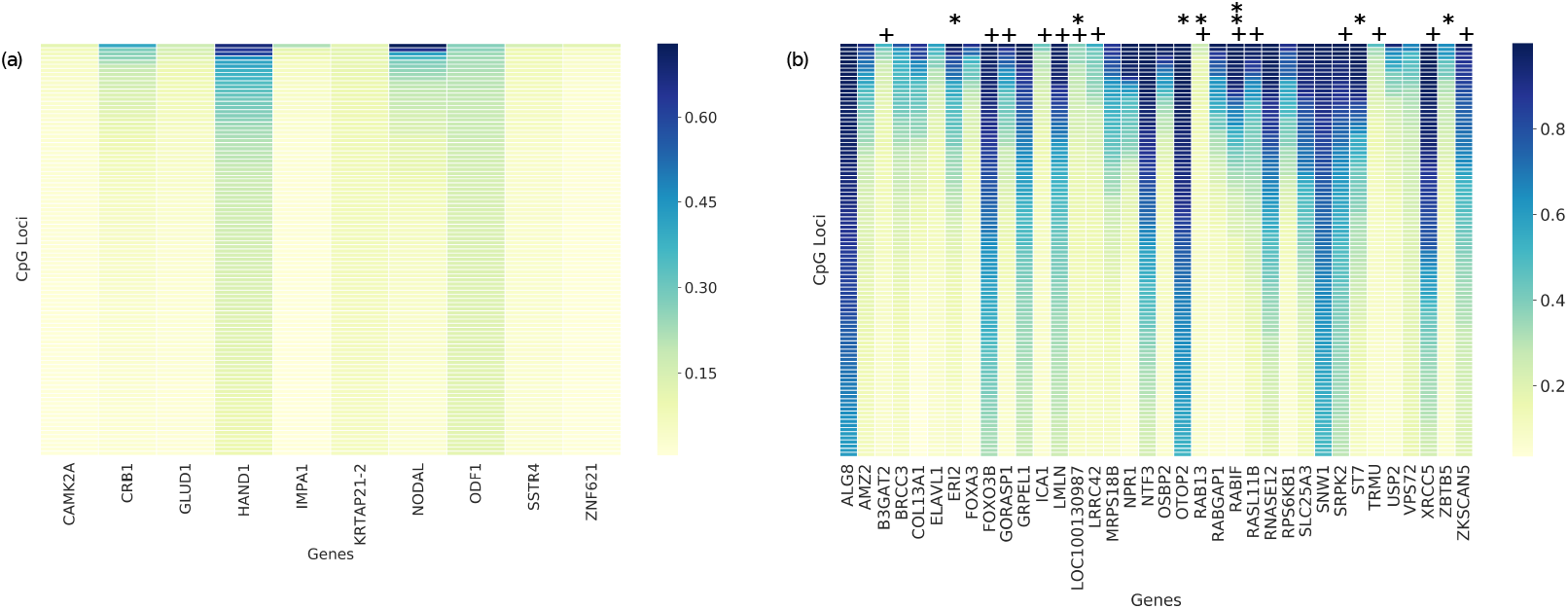
Attention probabilities per gene and CpG. Columns are genes and rows are CpGs, where for each gene the CpGs were sorted from high to low attention score and the first 100 CpGs are depicted (note that any specific row corresponds to a rank in the attention vectors and not to a specific CpG). (a) Sequence-based attention - genes with at least one attention score > 0:1. (b) Distance-based attention - genes with a median attention score > 0:1.

The most notable gene is HAND1. As mentioned in Section III-B, HAND1 is a transcription factor that plays a key role in the differentiation of cell lineages during human embryonic development. To assess the possible relationship between HAND1 and DNA methylation in our dataset, we retrieved the sequences surrounding the top CpGs for HAND1 (recall that attention is determined based on the CpG’s ambient sequence, in this case), and tested whether they are significantly enriched with the HAND1 binding site motif - GTCTGG [20], [16] as compared to the lower ranked CpGs. Such a case would indicate that the CpGs proximal to this motif produce higher attention scores for HAND1, linking their methylation level to HAND1 binding. We performed this analysis using DRIMUST [25], a tool which analyzes motifs enriched at the top of ranked lists. We took the top 20 CpGs for HAND1, with an average attention value of 0.32, along with the bottom 20 CpGs, with an average attention value of 0.07, and inserted them in ranked order. The top significantly enriched motif - GTCTGA - was indeed nearly identical to the known HAND1 binding motif, with p-value < 3*e*^−07^, as can be seen in Figure 5 (a). Furthermore, only the top 18 CpGs contained this motif and the top 10 contained it more than once. None of the bottom 20 sequences contained this motif. To the best of our knowledge, the HAND1 gene has not been previously associated directly with the process of methylation at or near HAND1 binding sites, yet our findings show that this may be the case.

Another prominent gene, which was given a high attention score by multiple CpGs, is NODAL. Interestingly, NODAL is also associated with embryogenesis [38]. In fact, in adult tissues it is not normally activated, except in the case of unhealthy tissues, and specifically cancer tissues, in which it known to re-express [39]. NODAL plays a crucial part in the Nodal Signaling Pathway (NSP). In this signaling pathway, NODAL is responsible for instigating the transcription of multiple target genes, which is likely part of the reason why the model attributed NODAL with a high attention value for multiple CpGs. According to a recent experiment conducted on mouse embryos, elevated NODAL levels may be linked with increased DNA methylation [9]. Combining this finding with the fact that our model attributes high attention values to NODAL across multiple CpGs when determining their methylation levels, indicates this may also be the case in humans, and specifically in cancer tissues. Performing motif analysis here, we discover a single significantly enriched motif - CGGCGGC (p-value < *e*^−10^) as seen in Figure 5 (b). Here too, the motif appears only in the top 20 sequences, and multiple times in the vast majority of them (the top 8 alone contain 37 occurrences, and all together the top 20 contain 82 occurrences).

**Fig. 5.**
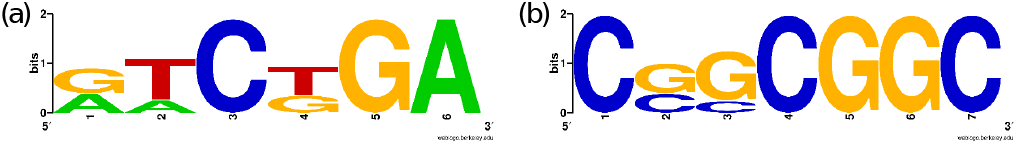
The most significantly enriched motifs in sequences with high sequence-based attention scores for HAND1 (a) and NODAL (b). The HAND1 motif is near-identical to its known binding site motif GTCTGG. The logos were generated by providing DRIMUST with a ranked list of 40 sequences - the ambient sequences of the top 20 CpGs with the highest sequence-based attention score, followed by those of the bottom 20. The logos were created using WebLogo [8] applied to the output of DRIMUST[25]

#### Distance-based attention

For the distance-based analysis, we obtained each gene's attention vector in a similar fashion to that described in the sequence-based section. In this case we fed the distance-based neural network with all unique distance vectors from the test set, resulting in one distance-based attention vector per CpG. We then sliced across CpGs to obtain all attention scores attributed to this gene (with one score provided by each CpG), and sorted its scores from high to low. The distance-based attention resulted in thousands of genes with at least one attention score larger than 0.1. Hence, for display purposes, we further refined the list of genes to include those for which the top 50 CpGs were larger than 0.1. This resulted in 36 genes for which the top 100 CpGs are shown in Figure 4 (b).

We hypothesized that the high attention scores may be explained by two main mechanisms: (1) in-cis effect: the CpGs reside within the 10,000 base-pair window for that gene and are therefore directly associated with it; or (2) in-trans effect: the gene enables the model to distinguish between different methylation profiles, and is therefore used by it to predict the methylation level for multiple CpGs in different genomic locations. We examine both options for every gene in Figure 4 (b). To assess whether the first hypothesis is true, we tested for a significant enrichment of nearby CpGs at the top of the attention list of each gene *g*. For this purpose, we took the gene’s sorted list of CpGs and labeled every CpG as 1 or 0 depending on the distance of *g* to that CpG (1 indicates ¡ 10K). This way the top of the list contains the 1/0 indicators corresponding to the CpGs with the highest attention. We then tested for statistical enrichment of 1s at the top of this list using the minimum hypergeometric (mHG) test [11]. This analysis yielded 7 genes for which the high attention values were significantly enriched with CpGs residing within the window limit (p-value < 0.05). These genes are marked with the asterisk symbol (*) in Figure 4 (b). Specifically, RABIF had a p-value < 0.003 and is marked with (**).

To further explain the second hypothesis, we will use FOXO3B as a simple example. Notice that FOXO3B is not marked with an asterisk, implying that its high attention scores are not associated to nearby CpGs. Instead, the model might have learned that a high expression level of FOXO3B is associated with positive methylation of a certain set of CpGs. During training, such a relationship is easily learned through the distance vector (the input to the distance-based attention mechanism). The distance vector provides the model with a means by which to identify the CpG, so that the next time the model encounters its distance vector, it will attend to FOXO3B (in our example). Recall, however, that the test set does not contain previously seen CpGs. Hence, the model cannot identify a test CpG based on its distance vector, unless it closely resembles one already seen in the training data. Therefore, given a new CpG from the test set, the model would only be able to identify such a gene-CpG relationship based on the CpG's proximity to a previously seen CpG (methylation levels of neighboring CpGs are closely linked [44]). To test in-trans roles for the genes in Figure 4 (b), we use the same approach as above, labeling 1 if the CpG has an adjacent CpG (in the ordered list) with the same distance vector and 0 otherwise. This resulted in 14 genes with 1s at the top (p-value < 0.003), marked with (+) in Figure 4 (b). While the existence of neighboring CpGs in the test set does not guarantee that the training data contains a neighboring CpG from which to learn, it indicates of a higher likelihood that one such CpG exists (recall that the data was randomly partitioned into training, validation and test). Looking at the locations of the top 20 CpGs for FOXO3B (p-value ¡ 0.0001), as seen in Supplementary Figure S1, we can see these CpGs reside in 8 different chromosomes and many of them have a nearby CpG. FOXO3B is a member of the forkhead family of DNA binding proteins, which is consistent with this observation [5].

Curiously, there are two prominent genes that do not fit either of the models above - namely, ALG8 and SNW1, both of which have a high attention score across at least 100 CpGs. This indicates that the model found them to be overall relevant, i.e. their expression level (whether on its own or combined with the expression levels of other genes) can help determine methylation levels for a large number of CpGs in many different genomic locations. These genes were studied in the context of methylation in breast cancer: ALG - [14], [33] and SNW1 - [36], [31].

### Motifs Learned by the CNN

In this section we looked into the representations learned by the CNN for motif detection. Specifically, we took each learned filter, and 0-1 scaled each row (representing a single nucleotide) to obtain the position probability matrix used for generating sequence logos. Figure 6 described six motifs detected by the CNN. One representation that stands out appears in the Figure 6 (b) and indicates the importance of both the individual CpGs that appear within the surrounding sequence, as well as the existence of multiple, consecutive combinations of Cs and Gs, most likely representing dense CpG occurrences - a hallmark of CpG islands.

Another prominent motif is the CA-motif in Figure 6 (a). This motif has been shown to modulate alternative splicing of mRNA [18], which is also thought to be regulated by methylation [27]. CpA repeats (TpG repeats) are also hallmarks of past methylation activity due to conversion of CpG to TpG when deamination follows methylation [7].

In another filter, we also identified the TATA motif (or TATA box), a core promoter element [43] seen in Figure 6 (c). Previous studies have shown that promoters residing in CpG-islands, and CG-dense regions in general, often lack this motif [43]. Hence, the model seems to distinguish between CpGs residing in CpG-dense regions and CpGs that are more isolated, specifically those residing in promoter regions that contain the TATA box.

The three remaining filters are likely related to the SP1 motif, which is especially known for its consensus sequence: GGGCGG and its reverse complement [34], clearly seen in Figure 6 (e). The SP1 motif is also known to have several consecutive Cs or Ts, which might be one of the reasons for observing Figure 6 (d) [32]. Figure 6 (f) contains DNA repeats which have also been linked to methylation [12].

**Fig. 6.**
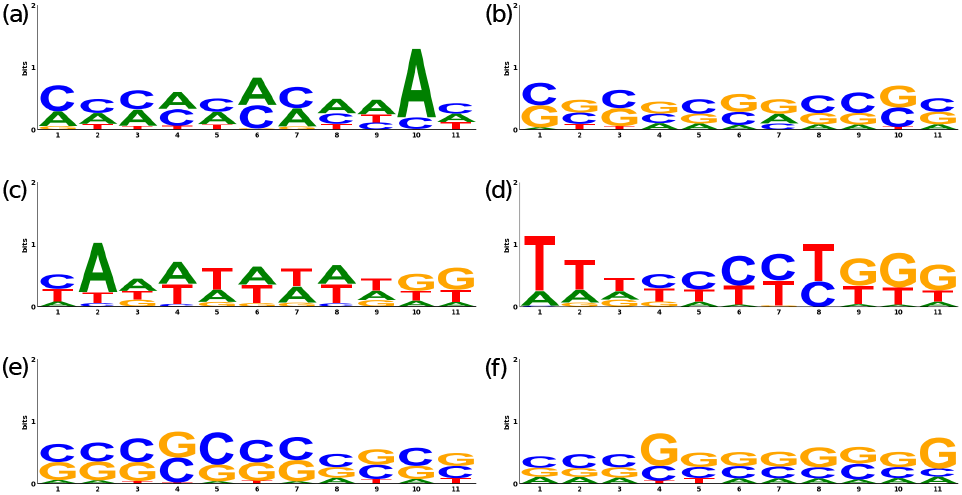
Six CNN filters and their learned weights presented as sequence logos. Top row: (a) The CA-repeat motif - a hallmark of methylation activity [30]. (b) The CpG dinucleotide and CpG-dense regions. (c) The TATA motif (TATA box) - a core promoter element [43]. (d), (e) The SP1 motif, known for having several consecutive Cs or Ts [32] and especially known for the consensus sequence: GGGCGG and its reverse complement [34], clearly observed in (e). (f) DNA repeats, also linked to methylation [12].

## V. Conclusions

DNA methylation is strongly related to disease development, and is therefore the focus of much research. Models that provide methylation predictions could speed up future research and improve our understanding on how epigenetics may be involved in physiopathology. In this paper, we provided a general model that supplies such predictions based on the ambient sequence at the CpG of interest and the sample’s gene expression profile. Our model is comprised of three sub-models that enable us to provide more accurate predictions under certain gene-CpG proximity conditions. We demonstrated the model’s capability of generalizing across both CpGs and samples by testing on completely separate sets of CpGs and subjects. Our model is highly interpretable, avoids incorporating prior knowledge, provides continuous predictions and is not limited to any subset of CpGs, thus improving upon previous models.

Furthermore, we demonstrate the power of using an attention mechanism on gene-expression data by analyzing its learned representations. Specifically, this enabled us to link HAND1 and NODAL to methylation activity. Our attention-based model, along with its analysis, provide a novel framework for future research that seeks to combine gene-expression data with genomic sequences and extract valuable insights from both. This framework could also be extended beyond gene-expression data, to include other genomic measurements.

## Supporting information

## Acknowledgment

We would like to thank the Yakhini Group, and specifically Leon Anavy and Oz Solomon, for valuable discussions and suggestions.

## Footnotes

1 https://github.com/YakhiniGroup/Methylation

