## Supplementary figures and images for "Predicting Methylation from Sequence and Gene Expression Using Deep Learning with Attention"

### Supplementary file 1

#### CpG Positions FOXO3B (top 20)

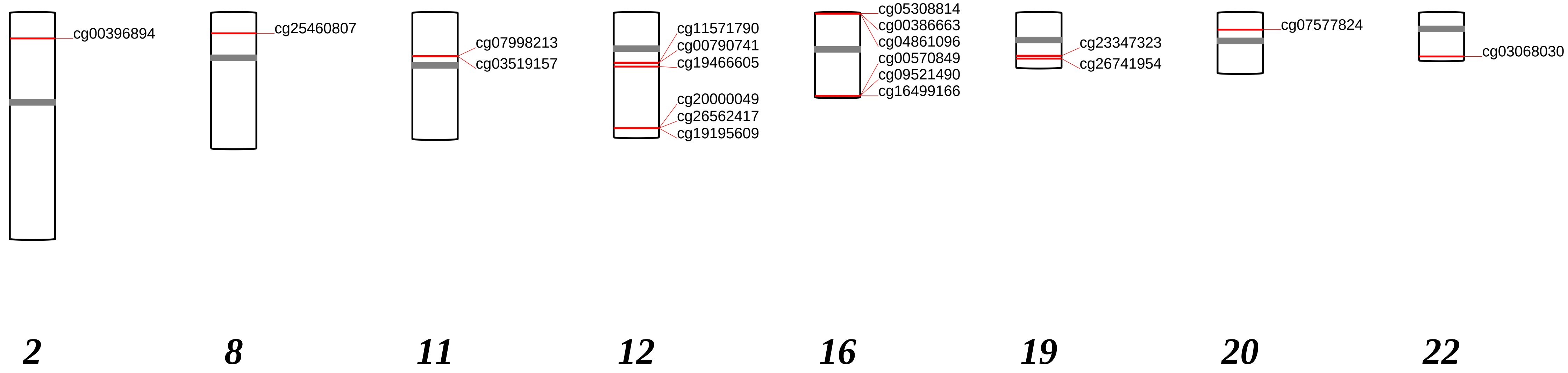
